# NEK1 modulates neurite outgrowth in motor neurons through coordinating retromer formation

**DOI:** 10.1101/2024.05.06.592854

**Authors:** Xuan Huang, Juan A. Oses-Prieto, Riki Kawaguchi, Laura Perrault, Devlin Frost, Kuchuan Chen, Feng Guo, Bingjie Zhang, Pinghan Zhao, Tammy Szu-Yu Ho, Roshan Pandey, Daniel Geschwind, Alma L. Burlingame, Clifford J. Woolf

## Abstract

Loss of function of a cell cycle-associated gene *NEK1* causes amyotrophic lateral sclerosis (ALS), but how this leads to motor neuron degeneration is unknown. We studied the function of *NEK1* in human stem cell-derived motor neurons and found that loss of *NEK1* causes decreased neurite length accompanied by transcriptional alterations. We also found that NEK1 interacts with and modulates the formation of the retromer, and that impaired retromer function contributes to neurite outgrowth deficits. We identified SMC3, which interacts with NEK1 during the cell cycle, as a kinase substrate of NEK1 in motor neurons. Notably, loss of SMC3 not only recapitulates the decreased neurite outgrowth, but also affects retromer formation. We suggest that NEK1 interacts with multiple proteins in postmitotic neurons to coordinate retromer formation, and that loss of this leads to impaired neurite outgrowth.

## INTRODUCTION

Amyotrophic lateral sclerosis (ALS) is a devastating neurodegenerative disorder with a complex pathology^1^. Diverse genetic mutations have been identified in patients that cause degeneration of motor neurons. While the impact of these mutations on different cell types contributes to ALS disease progression, their toxicity in motor neurons is a major driver of ALS via diverse mechanisms, including impaired proteostasis, disrupted RNA processing, and cytoskeletal and transport deficits^2,3^. Studies of ALS-associated genes have expanded our knowledge of ALS pathogenesis and revealed novel aspects of how the healthy nervous system works, but for many recently identified ALS-associated genes, their functions remain unknown.

NIMA-related kinase 1, NEK1, belongs to the NIMA (never in mitosis A) kinase family which play key roles in regulating different cell cycle events^4^. Loss-of-function (LOF) mutations in *NEK1* cause ALS, accounting for 3% of European and European-American patients^5–7^, but how mutations in a cell cycle-associated gene lead to ALS is not understood. NEK1 participates in multiple cellular events during the cell cycle progression^8,9,10^ including the cohesin complex removal^11,12^. NEK1 also regulates other functions beyond the cell cycle, such as mitochondrial permeability^13^ and cilia formation^14^ and is associated with both polycystic kidney disease^15^ and axial spondylometaphyseal dysplasia^16^. Nevertheless, the precise molecular machinery through which NEK1 exerts these diverse functions is unclear, and how NEK1 dysfunction affects the nervous system and contributes to ALS pathogenesis needs to be further investigated.

Impairment of the retromer, a multimeric protein complex involved in endosomal trafficking pathways, has been implicated in neurodegeneration. The retromer regulates protein recycling from the endosome and deficits in its function disrupts protein sorting and degradation ^17–19^. Mutations in a retromer complex subunit VPS35 cause Parkinson’s disease^20^, and loss of VPS35 decrease the neurite transport of Amyloid precursor protein^21^. While it was recently reported that NEK1 regulates retromer function in mouse fibroblasts and impacts proteostasis^22^, whether retromer abnormalities play a role in ALS pathogenesis is not clear.

To understand the function of *NEK1* in motor neurons, and to explore whether and how a loss of *NEK1* function in these neurons contributes to ALS, we examined the role of *NEK1* in stem cell-derived human motor neurons. Stem cell-derived neurons are a powerful model to study neurological disorders of genetic origin, and many ALS hallmarks have been recapitulated in patient-derived neurons, including decreased neurite length^23–25^. We found that loss of *NEK1* in motor neurons leads to decreased neurite length, increased ER stress, increased cell death, and is accompanied by altered expression of multiple neurite outgrowth and protein transport related genes. We further identified that NEK1 interacts with VPS26B, SMC3, and microtubules to modulate neurite outgrowth and retromer formation. We propose that NEK1 orchestrates the formation of the retromer in human motor neurons, and that loss of this function leads to impaired neurite outgrowth, which may lead to neurodegeneration.

## RESULTS

### Motor neuron NEK1 is located in the cytoplasm and is induced by impaired protein transport

To study the function of NEK1 in postmitotic neurons, we generated motor neurons from human stem cells using a dual-SMAD inhibitor mediated differentiation protocol and this neuronal population was isolated using magnetic associated cell sorting (MACS) with an anti-NCAM antibody at the end of the differentiation^26^ (Fig 1A). After the MACS >90% of the cells were Tuj1^+^ neurons and ∼50% were Isl1^+^Tuj1^+^ motor neurons (Fig S1A, S1B). While it is well established that NEK1 participates in multiple processes during the cell cycle^10,11^ we found that NEK1 is present in these postmitotic motor neurons. Furthermore, NEK1 mRNA and protein levels were higher in neurons compared to stem cells (Fig 1A, 1B), suggesting that NEK1 has cell-division-independent functions in these neurons.

**Figure 1:**
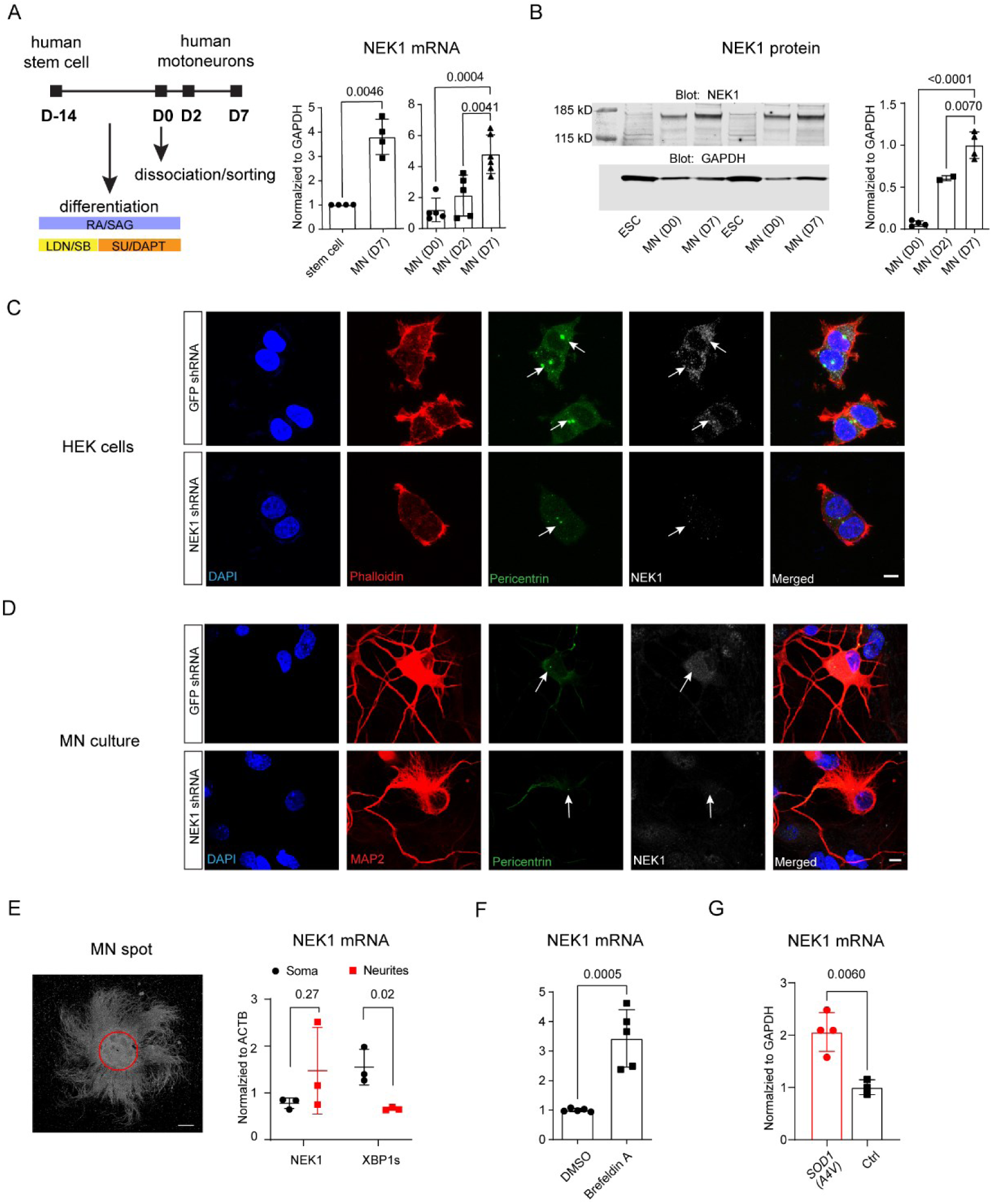
NEK1 is expressed in the cytoplasm of motor neurons and is upregulated during neurite outgrowth and induced by stress. A. qRT-PCR analysis to compare NEK1 transcript levels in human stem cells and differentiated motor neurons (MN) (n=4), (DIV=0, 2, and 7) (n=4∼6). Motor neurons are differentiated *in vitro* for 14 days using small molecules, dissociated/purified at DIV=0, and cultured for up to one week (DIV=7). B. Western blot analysis of NEK1 protein in human stem cells and differentiated motor neurons (DIV=2 and 7) (n = 2∼4) C. Representative confocal images of HEK cells treated with control or NEK1 shRNA, stained for the centrosome (pericentrin) and NEK1, and counterstained with DAPI and phalloidin. Scale bar = 10 µm D. Representative confocal images of 3-week-old motor neurons treated with control or NEK1 shRNA, stained for MAP2, centrosome (pericentrin) and NEK1, and counterstained with DAPI. Scale bar = 10 µm E. qRT-PCR analysis of NEK1 transcript expression in soma and neurite area collected from 2-week-old motor neuron spot cultures (n=3) (Scale bar = 800 nm) F. qRT-PCR analysis of NEK1 transcript in motor neurons (DIV=4∼7) treated by brefeldin A (2-10 µM, overnight) (n=5) G. qRT-PCR analysis of NEK1 transcript in motor neurons (DIV=7) derived from a human patient stem cell line bearing the *SOD1(A4V)* mutation, and the isogenic control where the mutation is corrected (n=4 and 3 respectively)

NEK1 is recruited to the nucleus, as a part of the DNA damage response^27^, and to the centrosome for spindle apparatus and cilia formation^28^. To understand the function of NEK1, we compared its expression in dividing HEK cells and postmitotic motor neurons. In HEK cells NEK1 is co-localized with centrosome puncta (pericentrin) and also diffusely expressed in the cytoplasm (Fig 1C). However, while NEK1 nuclear puncta were occasionally observed in motor neurons (Fig S1C), we mostly found a diffuse cytoplasmic localization which did not colocalize with the neuronal centrosome (Fig 1D). A diffuse expression pattern in both the neuronal soma and in neurites was also observed for exogenously expressed human NEK1 (Fig S1D). To better identify the cellular localization of NEK1, we seeded a high density of motor neurons as a spot culture from which axons grow outward in a radial fashion allowing us to collect proteins in either the soma or neurites (Fig 1E). As expected, XBP1s, a nucleus-located transcription factor, was enriched in the soma, however, NEK1 mRNA could be detected in both (Fig 1E), indicating a potential role for NEK1 in axonal function.

Members of the NIMA family participate in different stress responses and exhibit a stress-induced expression^27,29^. In human motor neurons brefeldin A, a molecule that blocks protein transport, increased NEK1 transcript levels (Fig 1F). In contrast, thapsigargin and 2-deoxyglucose, that interfere with ER calcium homeostasis and mitochondrial function respectively, had no effect on NEK1 levels, suggesting that it’s only a stress specific to protein transport that induces NEK1 upregulation (Fig S1E). ALS-associated mutations in SOD1 inhibit protein transport from the ER^30^. We found that NEK1 expression was upregulated in patient-derived motor neurons bearing the *SOD1(A4V)* mutation, further implicating a possible link between NEK1 and protein transport (Fig 1G). Overexpression of NEK1 in wildtype motor neurons increased neuronal survival (Fig S1F) suggesting that the upregulation of NEK1 could be a neuronal protective response to the stress induced by impaired protein transport.

### Loss of NEK1 in human motor neurons decreases neurite length

To understand the association of *NEK1* with ALS neurodegeneration we next studied the effects of a loss of *NEK1* in human motor neurons. We generated a homozygous and a heterozygous knockout stem cell line from WAO1 embryonic stem cells, using guide RNAs targeting different genomic areas (Fig 2A, S2A). NEK1 mRNA and protein levels were decreased to different extents in the knockout stem cells (Fig S2B, S2C) and in the motor neurons differentiated from them (Fig 2B), without affecting the percentage of differentiated motor neuron (%Isl1^+^Tuji1^+^/Tuji1^+^) (Fig S2D).

**Figure 2.**
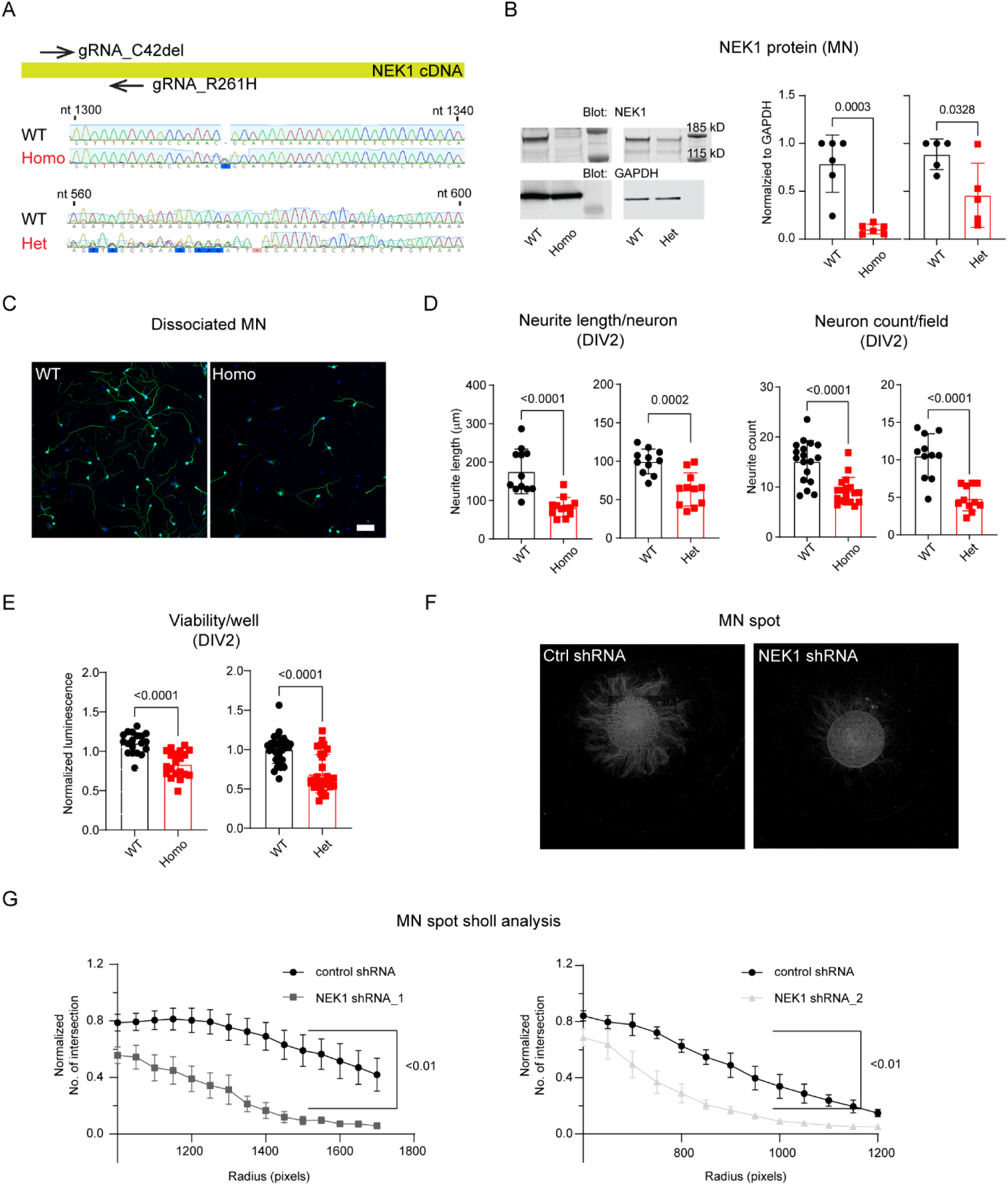
Loss of NEK1 leads to decreased neurite length. A. DNA chromatogram showing homozygous (homo) and heterozygous (het) NEK1 knockout stem cells and their wildtype (WT) controls after CRISPR gene editing B. Western blot analysis of NEK1 protein in homozygous (n=6) and heterozygous knockout motor neurons collected at DIV7 (n = 5) C. Representative epifluorescence images from wildtype (WT) and homozygous knockout (Homo) motor neurons at DIV2 (Scale bar = 100 µm) D. Quantification of neurite length per neuron and neuron count per well for both homozygous knockout and heterozygous knockout (Het) and their controls (n=11-18) at DIV2 E. Viability measurement of homozygous and heterozygous knockout motor neurons and their controls at DIV2 by the CellTiterGlo assay (n=19) F. Representative images of DIV8 motor neuron spot cultures treated with control or NEK1 shRNA G. Neurite outgrowth quantification of motor neuron spot culture by Sholl analysis (n=4-6)

ALS can be caused by multiple distinct genetic mutations affecting several different downstream pathways, amongst which is axon degeneration, a common feature shared among several different ALS models^1^. Terminal degeneration of axons innervating the neuromuscular junction is one of the earliest manifestations of ALS^31^, which manifests as neurite outgrowth defects *in vitro*^23,25^. To study whether an absence of NEK1 had any effect on axons we traced neurites from individual neurons two days after sorting and replating (DIV2) and found that neurite length was significantly reduced in motor neurons differentiated from both homozygous and heterozygous NEK1 knockout stem cells (Fig 2C, 2D, S2E). The reduced neurite length was accompanied with a reduced neuron count (Fig 2D) reflecting decreased motor neuron viability (Fig 2E). However, the decreased neurite length was not caused by the reduced neuron count (Fig S2F), and the diminished growth could be rescued by blebbistatin (Fig S2G), a small molecule that promotes axon re-growth and substrate adhesion^32^. The decreased neurite length could not be rescued by the apoptosis inhibitor ZVAD or the necroptosis inhibitor necrostatin-1 (Fig S2H), suggesting that loss of *NEK1* specifically causes an early impairment of neurite outgrowth which manifests at DIV2. To verify that the observed phenotype was not caused by gene editing artifacts or potential NEK1 function during differentiation, we also knocked down NEK1 expression in differentiated postmitotic neurons by shRNA and examined axon outgrowth in spot cultures. shRNA constructs targeting NEK1 in the post-differentiation motor neurons significantly also reduced axon length (Fig 2F, 2G).

### Loss of NEK1 in human motor neurons leads to increased ER stress and cell death, as well as altered excitability

Axon degeneration in ALS presents early and then typically progresses to neuronal loss^1^. To study whether loss of *NEK1* leads to other ALS phenotypes, we cultured the motor neurons longer than two days, the first time point when we first observed a decreased neurite length. After culturing for one week, we found increased ER stress responses (Fig 3A), cell death (Fig 3B), and caspase activity (Fig 3C) in homozygous NEK1 knockout motor neurons. We also co-cultured human motor neurons with mouse glia for 4 weeks (Fig S3A) to promote neuronal maturation^26^ and found increased neuronal firing as well as increased capacitance in the homozygous knockout neurons, indicating altered membrane excitability (Fig 3D). A similar but less dramatic change of membrane properties was found in heterozygous knockout motor neurons (Fig 3E). While a decrease in neurite length was the earliest pathological phenotype observed, loss of *NEK1* eventually led to dysfunction of multiple pathways, much like several other ALS models^33^.

**Figure 3.**
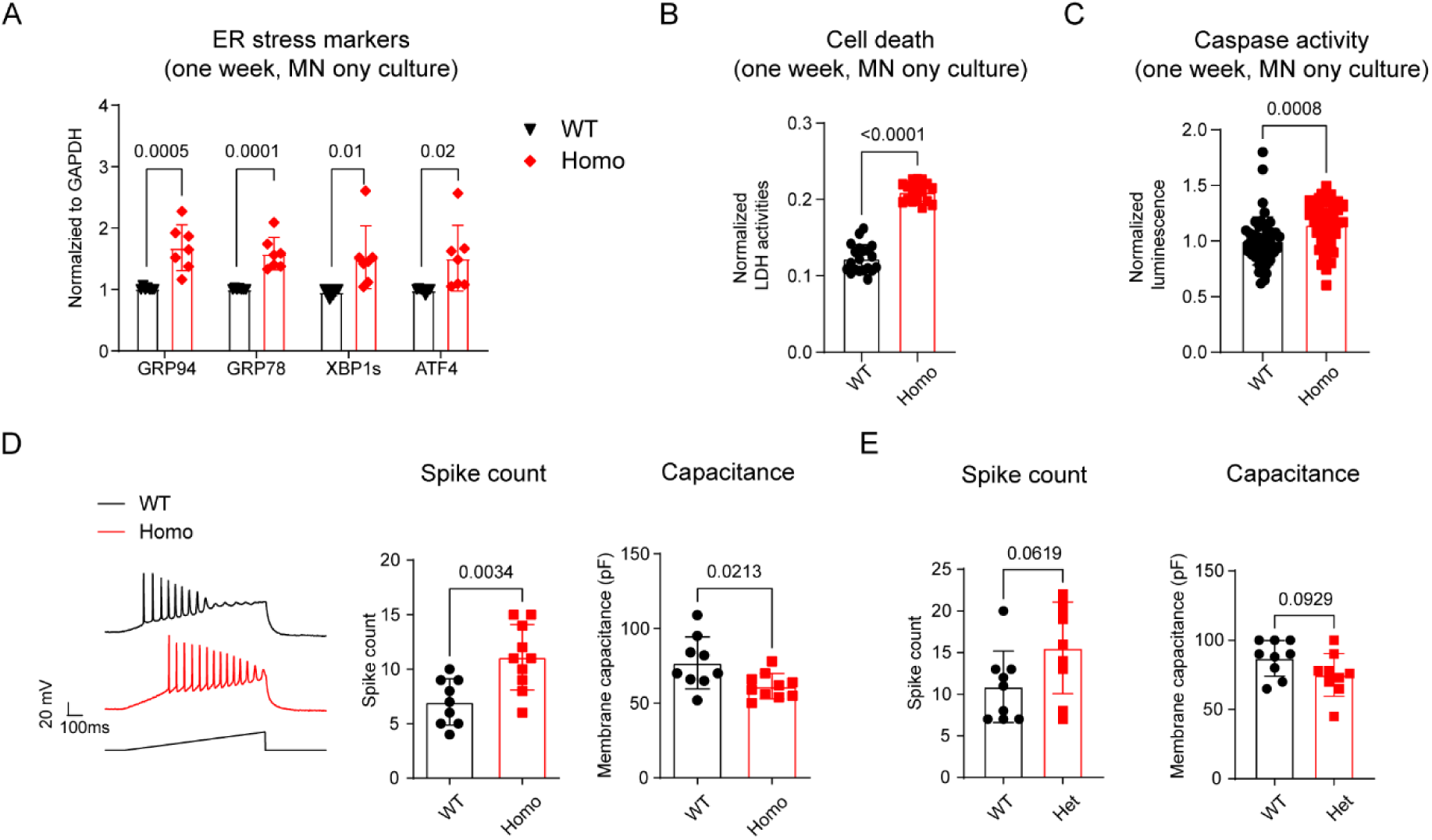
Loss of NEK1 leads to increased ER stress, cell death, and neuronal excitability in human motor neurons. A. qRT-PCR analysis of ER stress markers expressed in one-week-old wildtype and homozygous knockout motor neurons (n=7) B. Measurements of cell death in one-week-old wildtype and homozygous knockout motor neurons (n=20) C. Caspase 3/7 activity in one-week-old wildtype and homozygous knockout motor neurons (n=50) D. Representative images of wildtype and homozygous knockout motor neuron firing in response to a 200pA current injection, and measurements of the number of spikes and membrane capacitance (n=9-10) E. Measurements of MN firing properties in wildtype and heterozygous knockout motoneurons cocultured with glia (n=9)

Interestingly, we found that co-culture with glia prevented the increase in ER stress (Fig S3B) and cell death (Fig S3C) seen in neurons cultured without glia. Glia support neuronal health, and their dysfunction contributes to ALS progression^34^. Our observations indicate that glia are involved in the pathogenesis induced by loss of *NEK1* and suggest that an additional stressor may be required for the early axon degeneration to progress to other cellular dysfunctions.

### Impaired retromer formation in NEK1 knockouts contributes to decreased neurite outgrowth

To characterize the molecular mechanisms underlying the early decreased neurite outgrowth resulting from loss of *NEK1* in motor neurons, we profiled the transcriptome at DIV2, when a significant neurite length reduction but no obvious cell death was present in NEK1 knockouts. A major alteration in the motor neuron transcriptome was identified with homozygous knockout (Fig S4A, Supplementary table 1). The most significant differential expressed genes (DEG) were linked to biological processes like morphogenesis, and cellular components include axons and synapses (Fig S4B, S4C), including ARHGAP22, MAP7D2, MSN, and STMN2 (Fig 4A). Notably, the expression of several cell cycle related genes, including CCND1, CDK1, and CDC6 were also increased (Fig 4B), as observed in Alzheimer’s disease patient-derived neurons^35^, suggesting a potential shared mechanism with other neurodegenerative conditions. In contrast, while ER stress markers were upregulated at DIV7, their transcript levels were not affected at DIV2, confirming a progression of different disease phenotypes with time (Fig 4C).

**Figure 4.**
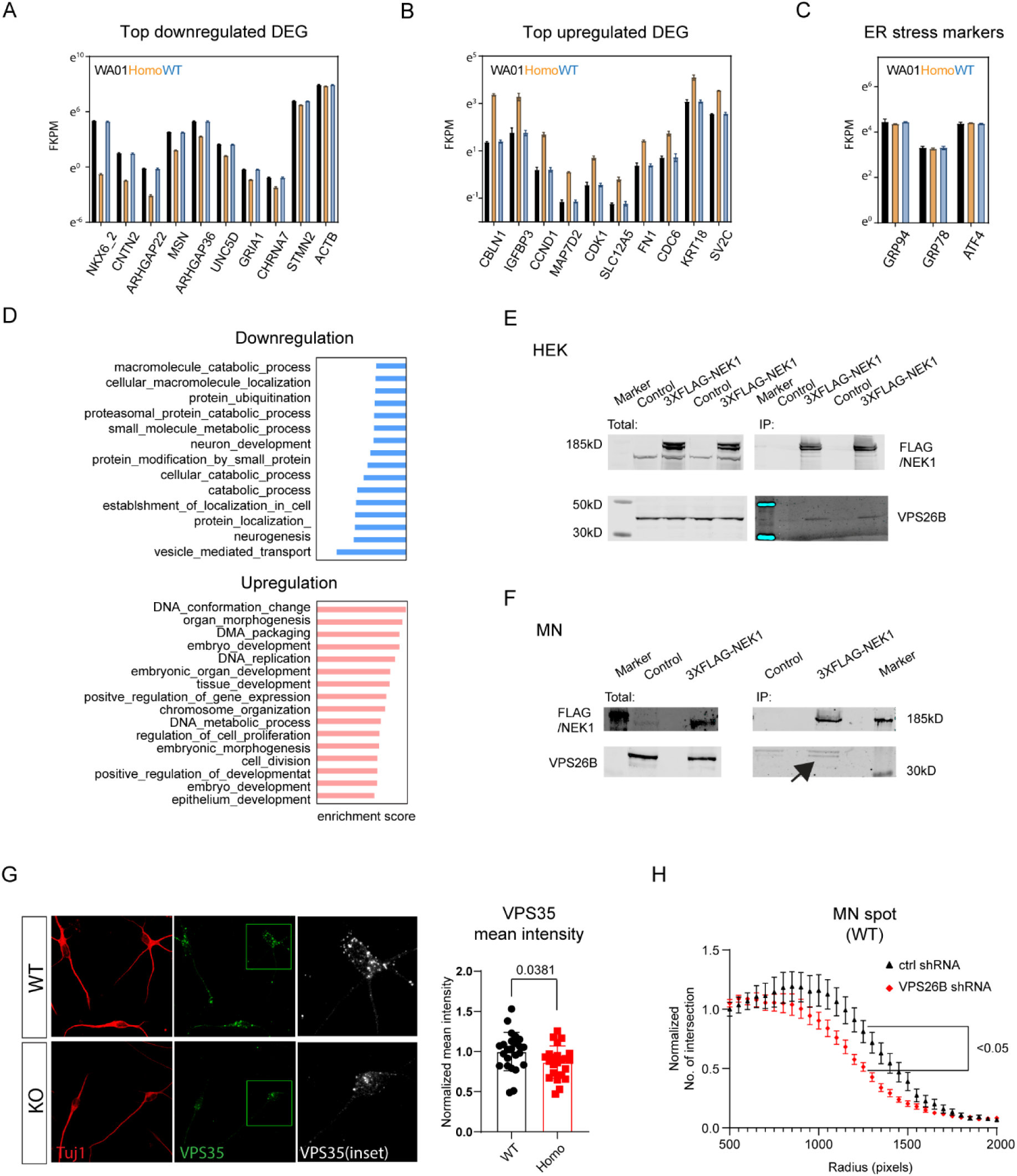
Impaired retromer formation contributes to decreased neurite outgrowth in NEK1 knockout motor neurons. A. Examples of most significantly downregulated transcripts in homozygous knockout MN (DIV2) B. Examples of most significantly upregulated transcripts in homozygous knockout MN (DIV2) C. Expression levels of ER stress transcripts (DIV2) D. Top down- and up-regulated gene sets in homozygous knockout MN revealed by GSEA E. Representative Western Blot images showing immunoprecipitation of VPS26B with NEK1 in HEK cells. HEK cells are transfected with the 3xFLAG-NEK1 construct or control construct then the protein is collected four days after transfection. F. Representative Western Blot images showing immunoprecipitation of VPS26B with NEK1 in motor neurons. Neurons were infected with lentivirus expressing the hsyn-3XFLAG-NEK1 construct or the control lentivirus expressing hsyn-EGFP at DIV0, and the protein was collected at DIV7. The specific band of VPS26B is identified by an arrow. G. Representative confocal image of wildtype and homozygous knockout MN (DIV2) stained for Tuj1 and VPS35, and the mean VPS35 intensity in neuronal soma is quantified (n= 29 and 23 cells respectively). H. Neurite outgrowth Sholl analysis of wildtype MN spots treated with control or VPS26B shRNA (n=8)

To explore whether dysregulation of the top DEG contributed to the decreased neurite length in NEK1 knock outs, we knocked down ARHGAP22, which encodes a Rho GTPase-activating protein regulating actin function^36^ which was downregulated in both homozygous and heterozygous NEK1 knockout motor neurons, as well as in homozygous knockout motor neurons generated from a different stem cell background (Fig S4D, S4E). However, knockdown of ARHGAP22 did not decrease neurite outgrowth (Fig S4F), indicating that dysregulation of this transcript accompanied the decreased neurite length, but was not its cause.

To identify potential upstream mechanisms leading to the decreased neurite length and dysregulated gene expression, we performed a gene set enrichment analysis (GSEA) to pin down altered pathways. Biological processes related to protein transport and catabolic processes were downregulated in the homozygous knockouts (Fig 4D), while establishment of protein localization was downregulated in the heterozygous knockouts (Fig S4G). NEK1 interacts with the retromer, a multimeric endosomal coat protein complex that regulates protein recycling^5,22^. We therefore asked whether NEK1 impacts neurite outgrowth through the retromer. Retromer subunits VPS26, VPS29, and VPS35 form a complex that recognizes recycling cargos^37^. We found that NEK1 interacts with VPS26B in both HEK cells (Fig 4E) and motor neurons (Fig 4F) and that loss of *NEK1* decreased retromer puncta intensities (Fig 4G, S4I). The retromer is pivotal for the glucose transporter GLUT1 to be recycled to the cell surface^38^. We found an accumulation of GLUT1 in cells lacking NEK1 (Fig S4J), confirming that retromer function is impaired by loss of NEK1. Furthermore, a loss of VPS26B in motor neurons not only decreased neurite outgrowth (Fig 4H) but also reduced expression of ACTB and ARHGAP22, two neurite outgrowth related transcripts also downregulated in *NEK1* knockout motor neurons (Fig S4H), suggesting that impaired retromer formation contributes to the decreased neurite outgrowth in *NEK1* knockout motor neurons.

### NEK1 interacts with and phosphorylates SMC3 in human motor neurons to regulate neurite outgrowth and retromer formation

To investigate how NEK1 affects retromer function, we overexpressed a kinase dead *NEK1(D128A)* mutant in homozygous *NEK1* knockout motor neurons and did not observe rescue of the decreased neurite length (Fig S5A), in contrast to overexpressing wildtype NEK1, indicating that the kinase activity of NEK1 is important for its neurite outgrowth function. Therefore, we set out to determine potential kinase substrates of NEK1 that may participate in neurite outgrowth control in motor neurons, and whether they influence retromer activity.

Stable Isotope Labeling with Amino Acids in Cell Culture (SILAC) is a metabolic labeling technique for mass spectrometric (MS)-based quantitative proteomics that typically requires near complete labelling of samples. While differentiation from dividing stem cells can promote metabolic label incorporation in the generated neurons, it is still not comparable to the incorporation in dividing cells. To overcome this constraint, we used isotopologues of Lysine for neutron-encoded stable isotope labeling^39^, which allows for SILAC analysis on samples with a low incorporation rate because the quantitation is done using only the labelled fraction of the proteome (Fig S5B). We got more than 50% labelling in 80% of the proteome, with the median labelling at 88% of each protein. We quantified more than 7000 phospho-peptides from four pairs of samples (two derived from WA01 WT:KO stem cells and two from another line SAH WT:KO), and more than 300 of them were significantly reduced (Fig 5A, Supplementary table 2). Among the most significantly downregulated phospho-peptides, two were derived from SMC3 and WASHC4, which are potential direct interactors with NEK1 (Fig 5A).

**Figure 5.**
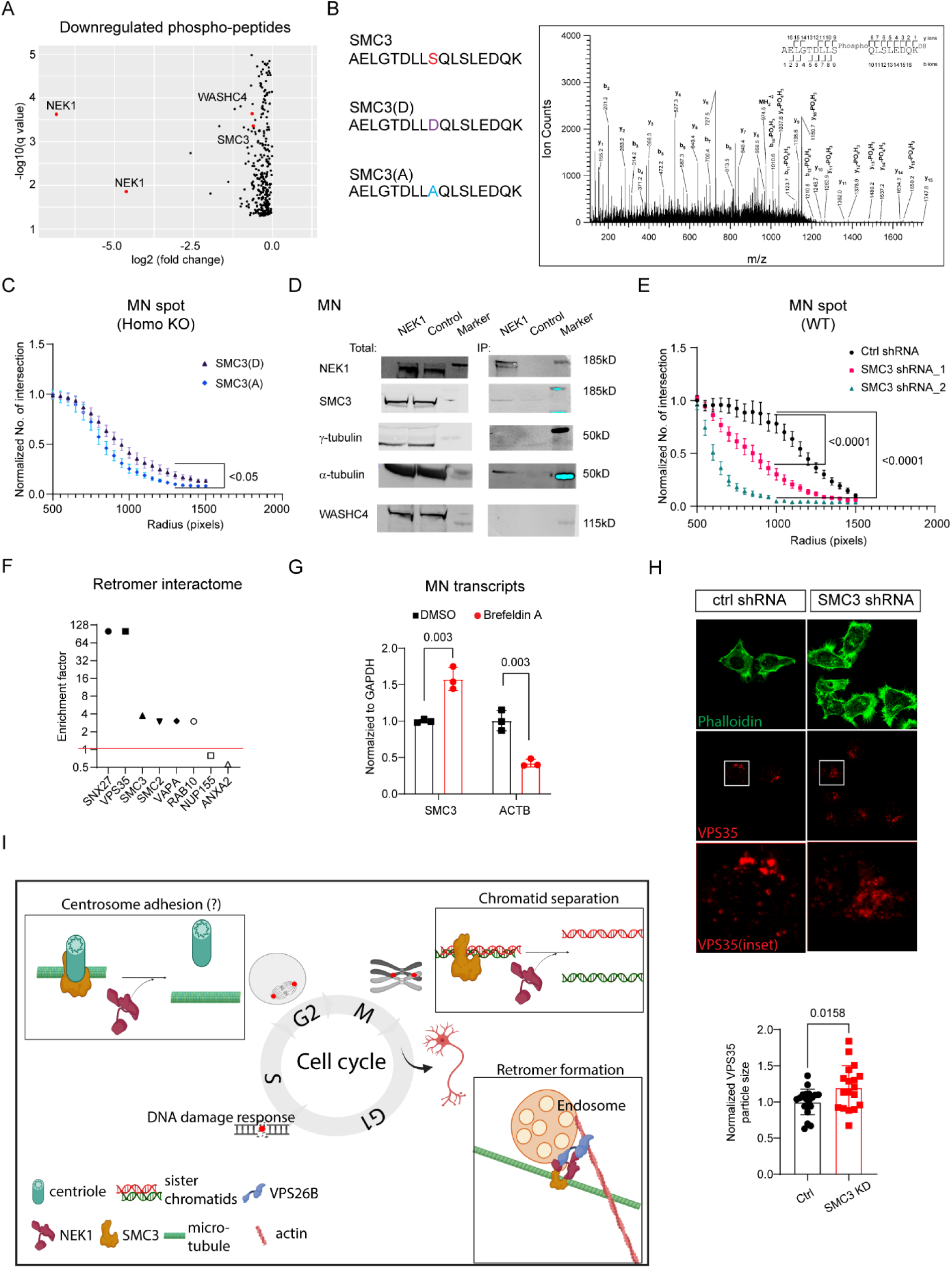
NEK1 interacts with and phosphorylates SMC3 in MNs to regulate neurite outgrowth and retromer formation. A. Volcano plot showing decreased phosphopeptides quantified from a Neucode-SILCA MS analysis B. Left: WT and mutated sequences (phosphomimic S to D, and non phosphorylatable S to A) around the S787 phosphorylation site in SMC3. Right: MS spectra of the SMC3 peptide. High-energy collision dissociation–tandem mass spectra obtained from precursor ion with mass 974.4916^+2^ found in LysC digests of D(8) (WT cells)/13C(6) 15N(2) (NEK1 KO cells)-Lysine labelled motor neurons, corresponding to the sequence A779 to K795 of human SMC3, phosphorylated at S787 and lysine D(8) labelled. Masses of the C terminal sequence ions (y) after y_9_ show an increase in mass of 80 Da over the unmodified sequence, pinpointing the location of the phosphate modified residue to S787. Observed sequence ions are labelled in the spectrum and marked in the sequence in the upper right corner. C. Neurite outgrowth Sholl analysis of homozygous knockout MN spot cultures overexpressing phospho-mimic SMC3(D) and phospho-silent SMC3(A) mutants (n=18-19) D. Representative Western Blot images showing immunoprecipitation of SMC3 and tubulin with NEK1 in motor neurons. Motor neurons are infected with lentivirus expressing the hysn-3XFLAG-NEK1 construct at DIV0, and the protein is collected at DIV7. E. Neurite outgrowth Sholl analysis of wildtype MN spots treated with control or SMC3 shRNA (n=8-19) F. Enrichment of SMC3 in retromer interactome. Adapted from published proteomics dataset quantifying proteins interacting with SNX27^42^. G. qRT-PCR analysis of SMC3 and ACTB transcript from wildtype motor neuron treated with brefeldin A (5 µM, overnight, n=3) H. Representative confocal image of SMC3 knockdown in Hela cells and its quantification. Cells were treated with control or SMC3 shRNA, stained for VPS35, counter stained with phalloidin-488, and quantified for the VPS35 particle size. Mean particle size of 21 and 18 images from three different experiments were compared. I. Proposed working model. During the cell cycle progression, NEK1-SMC3 interactions are involved in sister chromatid separation and may also affect centrosome adherence to microtubules. We propose that in postmitotic neurons, NEK1 interacts with VPS26B, SMC3, and microtubules, and that the NEK1-SMC3 interaction is important for creating the retromer tubule structure during the retromer formation (illustration generated by biorender.com).

SMC3 is a component of the cohesin complex that holds sister chromatids together during cell division, and it also contributes to microtubule-centrosome linkage^40^. WASHC4 is a component of the WASH complex, a nucleation-promoting factor (NPF) participating in tubule scission during retromer-mediated protein recycling^41^. To study whether SMC3 or WASHC4 are kinase substrates of NEK1, we generated phospho-mimic and phospho-silent mutants and evaluated whether a reduced phosphorylation of either WASHC4 or SMC3 contributed to the decreased neurite length in the NEK1 KO motor neurons. For the SMC3 peptide the phosphorylation site was unambiguously assigned at S787, and for WASHC4 there was not spectral information to resolve between T1156 or S1157 (See MS spectra and mutant sequences in Fig 5B and Fig S5C). Expression of the phospho-mimic mutation SMC3(D) increased neurite outgrowth in NEK1 knockout motor neurons compare to overexpressing phospho-silent SMC3(A) (Fig 5C), while phosphorylation of WASHC4 had little effect (Fig S5D). We then investigated their interaction by immunoprecipitation and found that in both HEK cells (Fig S5E) and differentiated human motor neurons (Fig 5D), NEK1 interacts with SMC3 but not WASHC4, suggesting that SMC3 but not WASHC4 is a kinase substrate of NEK1 in human motor neurons and contributes to the neurite outgrowth promotion by NEK1. Consistent with this, knockdown of SMC3 recapitulated the decreased neurite outgrowth of the NEK1 knockout (Fig 5E) while a loss of WASHC4 did not (Fig S5F).

Interactions between NEK1 and SMC3 play a vital role in chromosome segregation^11^ in dividing cells and regulate centrosome activities^28,40^. We therefore hypothesized that NEK1-SMC3 interactions modulate retromer activities and that an impairment of these activities may decrease neurite outgrowth. In the retromer interactome^42^ both SMC2 and SMC3 are enriched with VAPA and RAB10, molecules involved in protein recycling pathways (Fig 5F). SMC3 expression was also upregulated when protein transport was blocked in motor neurons (Fig 5G), similarly to NEK1 (Fig 1F). We therefore examined whether a loss of SMC3 impacts retromer activities. While VPS35 puncta in control cells had a crisp distinct morphology, those formed in SMC3 knockdown cells aggregated together, suggesting impaired retromer formation (Fig 5H) and implying that SMC3 regulates the retromer, like NEK1. SMC3 interacts with microtubules^43^ and we found that NEK1 also interacts with α-tubulin, in both HEK cells and motor neurons (Fig S5E, 5D). The formation of a retromer tubular structure from the endosome is an important step during protein recycling, and requires participation of microtubule motors^44^. Our data suggest that an interaction of NEK1 with SMC3 regulates this process.

We conclude that in motor neurons NEK1 interacts with both VPS26B and SMC3 to orchestrate retromer activity and that a loss of this function leads to decreased neurite growth, which may contribute to the pathogenesis of ALS (Fig 5I).

## DISCUSSION

We have examined the role of a newly identified ALS gene, *NEK1,* in human motor neurons. Our major finding is that a loss of *NEK1* disrupts multiple ALS-associated pathways, with in particular, an early loss of neurite outgrowth. We suggest that this is the consequence of its role in regulating retromer formation, through a network of multiple proteins including VPS26B, SMC3, and microtubule components (Fig 5I). Our study reveals how NEK1 acts in human neurons, through regulating retromer function, and that disruption of this function may contribute to ALS pathophysiology. By sorting ion channels, nutrient transporters and signaling receptors, the retromer complex participates in multiple diverse cellular events, including morphogenesis, synaptic plasticity, nutrient uptake, and apoptosis, and it is also implicated in neurodegeneration^18^. We find that NEK1 expression in motor neurons is upregulated when protein transport is impaired and that NEK1 interacts with the retromer subunit VPS26B. Furthermore, retromer function is reduced in the absence of NEK1, and this specifically leads to an impairment of neurite outgrowth. These data collectively point to a contribution of impaired retromer-mediated protein transport to ALS-related pathology in motor neurons lacking functional NEK1. Further work is required to establish exactly what aspect of retromer activity is impacted. One possibility is the regulation of retromer tubule structure formation from the endosome, which requires opposing forces generated by actin and microtubule motors^44^ (Fig 5I). In motor neurons NEK1 interacts directly with α-tubulin, suggesting that it may affect tubulin-retromer coordination during retromer tubule formation.

Our study also reveals new functions of cell cycle-associated genes in neurons. During the cell cycle, NEK1 directly interacts with the cohesin complex and modulates its function to regulate sister chromatids activities^11,45^. While we have uncovered a role of NEK1 in regulating neurite outgrowth through retromer function, we found this is achieved, at least in part, through an interaction with, and phosphorylation of the cohesion complex subunit SMC3. This suggests that NEK1 and SMC3 interactions have multiple functions in different locations, such as sister chromatid cohesion inside the nucleus, centroid-microtubule connection in the centrosome, and retromer formation in neuronal soma and neurites. NEK1 interacts with SMC3, α-tubulin, and γ-tubulin in HEK cells, but not with γ-tubulin in motor neurons, reflecting the different subcellular location and biological function of NEK1/SMC3 in these cells. A cell cycle-independent neuronal function of cell cycle-associated genes is not limited to NEK1 and SMC3. We observed increased expression of multiple cell cycle markers in NEK1 knockout motor neurons, which is also observed in neurons derived from Alzheimer’s patient iPSCs^35^. Cell cycle re-entering in neurons is linked to neuronal cell death^46^. However, a handful of cell cycle regulators execute diverse neuronal functions separate from the cell cycle^47^. Stromalin, another subunit of the cohesin complex, regulates the size of synaptic vesicles^48^ indicating a cell cycle independent role of the cohesion complex. Given our findings that both NEK1 and SMC3 display cell cycle independent functions in motor neurons, upregulation of these genes is likely to be an indication of their participation in other functions in non-dividing motor neurons. What exactly these cyclin proteins do in healthy neurons and how they contribute to neurodegeneration when mutated, are important questions that need to be answered.

While we focused here on NEK1’s function in the cytoplasm of motor neurons, we cannot rule out a role in the nucleus (Fig S1C). NEK1 is recruited to DNA damage sites^27^ and loss of NEK1 leads to elevated DNA damage during the cell cycle^9^. Although we did not find evidence of DNA damage in NEK1 knockout motor neurons (data not shown), it is possible that environmental stressors or aging may trigger this in the absence of full NEK1 function. Considering the diverse proteins that NEK1 interacts with, and that an absence of NEK1 in motor neurons leads to the dysregulation of multiple biological pathways, all the specific functions of NEK1 in healthy motor neurons now need to be explored. We focused here on cell autonomous functions of NEK1 in motor neurons, but found that the progression over time to ALS phenotypes in NEK1 knockout motor neurons appears to need a second hit, as coculture with healthy glia prevented the motor neurons from developing ER stress or progressing to cell death. Elaborating if and how a loss of NEK1 function in cell types like astrocytes and microglia, affects neuron health, may further contribute to a greater understanding of how ALS-associated NEK1 mutations lead to the disease.

Overall, our study then shows that a loss of *NEK1* in human motor neurons contributes to impaired retromer function and an early reduction in axon growth. We also reveal how cell cycle-associated protein networks control cell cycle-independent functions in postmitotic motor neurons and that the loss of these contributes to their vulnerability. A reduction in retromer function may be then another pathological mechanism that leads to ALS.

## ACKNOWLEDGEMENTS

This project was funded by grants from the Department of Defense W81XWH2010077 (C.J.W.) and the Dr. Miriam and Sheldon G Adelson Medical Research Foundation (C.J.W.; R.K.; D.G.; A.L.B). Support for the Lumos platform was provided by the UCSF Program for Breakthrough Biomedical Research (PBBR).

## AUTHOR CONTRIBUTIONS

XH, JAO, RK, and CJW designed the research; XH, JAO, RK, LP, DF, KC, FG, BZ, and PZ performed the experiments; XH, JAO, and RK analyzed the data; TSH and RP helped provide experimental materials; XH, DW, ALB, and CJW oversaw the project; XH, JAO, RK, and CJW wrote the manuscript, with the help from the other authors.

## SUPPLEMENTAL INFORMATION TITLES AND LEGENDS

*Supplementary table 1: significantly differentially expressed genes in NEK1 homozygous knockout motoneurons*

*Supplementary table 2: significantly reduced phosphor-peptides in NEK1 homozygous knockout motoneurons*

## METHODS

### Stem cell culture and Motoneuron differentiation

Human embryonic stem cell line WAO1 (H1) was purchased from WiCell and human induced pluripotent stem cell line SAH-0047 was provided by the Sahin laboratory, Boston Children’s Hospital, Boston MA. Stem cells were expanded on Matrigel (Corning) coated plates using StemFlex medium (Thermo Fisher) and human motor neurons were derived based on a modified protocol for adherent cells ^25^. Briefly, stem cells were dissociated to single cells by accutase (Stem Cell Technologies) and two days later the differentiation media (DMEM:F12/Neurobasal 1:1, supplemented with N2, B27, Non-essential-amino-acid and glutamax) was added. For the first 6 days of differentiation, the media was supplemented with SB431542 (10uM), LDN (0.1uM), retinoic acid (1uM), SAG (1uM), and switched to retinoic acid (1uM), SAG (1uM), SU5402 (4uM), and DAPT (5uM) for the next 8 days. By the end of differentiation, cell culture was dissociated by accutase, and enriched through magnetic sorting using the anti-NCAM antibody (BD Biosciences)^26^. Purified neurons were plated in PDL/laminin coated plates and maintained in motoneuron medium (Neurobasal supplemented with N2, B27, Non-essential-amino-acid and glutamax) with ascorbic acid (0.2 mg/ml) and neurotrophic factors GDNF, BDNF and CNTF (R&D) (10ng/mL), and half the medium was refreshed twice a week. For long term culture (more than one week), the remaining progenitors in the culture were removed by adding 2.5 µM 5-fluorodeoxyuridine (FdU) and 2.5 µM 5-ethynyl-2’-deoxyuridine (EdU, 10mM stock) for 24-48 hours.

### CRISPR gene editing

gRNA sequence was designed through Benchling (www.benchling.com) and cloned into pCas9 vectors^49^ or synthesized in the RNP format (Synthego). 48h after electroporation (P3 Primary Cell 4D-Nucleofector™ Kit L, Lonza), stem cells were dissociated, sorted, and plated as single cells. Colonies were picked up manually and the genomic sequence around the gRNA targeting site was amplified by PCR, followed by sanger sequencing.

### RNA extraction and RT-qPCR

RNA was extracted from motoneuron culture using RNeasy Micro Kit (Qiagen) and then cDNA was generated using Superscript Vilo Synthesis Kit (Invitrogen) following manufacturer’s manuals. qPCR was performed on Applied Biosystems 7500 machine (Life Technologies) using Fast SYBR Green Master Mix (Roche).

### Cloning of NEK1 cDNA and site directed mutagenesis

Motoneuron NEK1 was cloned from transcribed cDNA by nested PCR and cloned into a lentiviral FSGW backbone. The D128A mutations was further introduced by site directed mutagenesis and the full NEK1 cDNA sequence was confirmed by Sanger sequencing.

### Lentivirus transduction and cDNA transfection

Lentiviral vectors expressing a specific cDNA construct were either generated by in house subcloning into a FSGW backbone or purchased from VectorBuilder (https://en.vectorbuilder.com/). For shRNA knockdown, MISSION lentiviral vectors were purchased from Sigma and a lentiviral construct targeting the GFP gene using the same pLKO backbone was used as a control ^50^. Lentivirus was prepared and concentrated via Virus Core in Boston Children’s Hospital and added to cells on the day of plating or one day later, withdrawn after 24h. To achieve a lower expression efficiency in motoneurons or to deliver the constructs to HEK cells, lipofectamine 3000 (Thermo Fisher) was used following the manufacture’s protocol.

### Immunocytochemistry

For immunofluorescence imaging, cells were fixed with 4% PFA for 20-30 minutes, permeabilized and blocked with 1% blocking reagent (Roche 11096176001), 0.5% BSA, and 0.2% Triton-X in PBS for 1 hour. Cells were then incubated overnight at 4°C with primary antibody and for 1 hour at room temperature with secondary antibodies. Nuclei were stained with DAPI or Hoechst 33342. Images were acquired and analyzed by the ArrayscanTM XTI (Thermo Fisher), or acquired by Zeiss 700/Zeiss 710 confocal microscopy and analyzed by ImageJ.

### Viability, caspase activity, and cell death assays

Human MNs were plated at a density of 20,000 per well in a 384 well format and compounds were added the same day if needed. Cellular viability (ATP) was measured after 48 hours using Cell Titer Glo kit (Promega) and caspase activity was measured after one week using Caspase Glo kit (Promega). To measure the cell death, cell media was replaced at DIV4 and collected at DIV9 to measure the activities of released Lactate dehydrogenase using LDH assay kit (Pierce).

### Western Blot and Immunoprecipitation

Cell culture were lysed by RIPA buffer containing protease and phosphatase inhibitors (Roche) followed by sonication. Th protein concentration was further quantified by BCA kit, denatured in loading buffer (Invitrogen), separated by SDS-PAGE (Invitrogen), transferred to nitrocellulose membranes (Invitrogen) and probed with antibodies. The protein signal on the membrane was visualized and quantified using Image Studio (Licor).

To perform immunoprecipitation, 3xFLAG-tagged NEK1 construct (VectorBuilder) was expressed to HEK cells by transfection, or to human motoneurons by lentivirus transduction. 4 days to one week after expression, cell culture was lysed using modified RIPA buffer (50mM Tris (pH 7.4), 2mM EDTA, 1% NP40, 150mM NaCl), quantified, loaded into anti-FLAG® M2 Magnetic Beads (sigma), and incubated at 4 degree for 2 h using a rotator. After washing off the beads and antibodies, the FLAG interacting protein mix was eluted by 3xFLAG peptides (Sigma).

### Spot culture and Sholl analysis

Human MNs were resuspended at a density of 100,000 per μL in growth media. 2μL spots were placed in the center of each well in a completely dried PDL/Laminin coated 24 well plate and incubated in a humidified 37°C degree incubator for 20 minutes. Growth media was then slowly added around the spot culture, and 2.5 µM 5-fluorodeoxyuridine (FdU) and 2.5 µM 5-ethynyl-2’-deoxyuridine were added to the medium for 24-48 hours to deplete dividing non-neuronal cells.

Images of spot cultures were taken in the Incucyte® live-cell analysis system (Essen Bioscience) at a 4X magnification. Neurite outgrowth analysis was performed using the Sholl analysis plug-in in ImageJ. Briefly, the spot culture images were converted into the binary mode first. Then concentric rings at 50 pixels step were generated around the cell body region and the number of axons intersecting with each ring was counted. A higher intersection number suggested more neurites could reach to the concentric ring.

### Whole cell patch clamp

Human MNs were plated on PDL/laminin-coated coverslips with mouse glia and cultured for 4 weeks in MN medium. Whole cell patch-clamp recordings were performed at room temperature in external solution (145 mM NaCl, 5 mM KCl, 2 mM CaCl2, 1 mM MgCl2, 10 mM glucose, and 10 mM HEPES (pH 7.4)) with the recording electrodes (2-5 MΩ) filled with an internal solution (135 mM K-Gluconate, 10 mM KCl, 1 mM MgCl2, 5 mM EGTA, and 10 mM HEPES (pH 7.2)). To induce action potential firing, cells were held at -65 mV and a depolarizing current ramp (0 to 200 pA in one second) was applied. Only neurons displaying mature morphology were used. Data were low-pass filtered at 2 kHz, digitized at 20 kHz, and analyzed using the pCLAMP 10 software suite (Molecular Devices).

### RNA sequencing

RNA-sequencing libraries were prepared using the TruSeq Stranded RNA (100ng) with RiboZero Gold. Libraries were indexed and sequenced by HiSeq4000 with 75 bp paired-end reads and at least 73M reads were obtained for each sample.

Quality control (QC) is performed on base qualities and nucleotide composition of sequences, mismatch rate, mapping rate to the whole genome, repeats, chromosomes, key transcriptomic regions (exons, introns, UTRs, genes), insert sizes, AT/GC dropout, transcript coverage and GC bias to identify problems in library preparation or sequencing. Reads were aligned to the latest human reference genome (GRCh38) using the STAR spliced read aligner (ver 2.4.0). Average input read counts were 81.9M and average percentage of uniquely aligned reads were 90.6%. Total counts of read-fragments aligned to known gene regions within the human refSeq gene model annotation (GRCh38) are used as the basis for quantification of gene expression. Fragment counts were derived using HTSeq program (ver 0.6.0). Genes with minimum of 5 counts for at least one condition (all replicates) were selected and differentially expressed genes were determined by Bioconductor package EdgeR (ver 3.14.0). Scripts used in the RNA sequencing analyses are available at https://github.com/icnn/RNAseq-PIPELINE.git. Raw and processesd data were deposited within the Gene Expression Omnibus (GEO) repository (www.ncbi.nlm.nih.gov/geo), accession number (GSExxxxx). Gene Set Enrichment Analysis (GSEA) was conducted on sorted gene list from differential expression (-signed logFC x log10p-Val) using MsigDB (ver7.0).

Neucode-SILAC mass spectrometry analysis Wiltype and homozygous NEK1 knock out stem cells from either WAO1 or SAH background were dissociated by accutase and plated on Matrigel coated 10cm dishes in StemFlex media (Thermofisher). Two days after seeding, different small molecules were added to the MN differentiation media for two weeks, followed by MACS, and plated for two days. NeuCode L-lysine:2HCL(3,3,4,4,5,5,6,6-D8) or L-Lysine:2HCL(13C6; 15N2) (Cambridge Isotope Laboratories) were used to label either wildtype or knockout cells. Lysine and Arginine deficient DMEM/F12 media (Thermofisher) and Lysine, Arginine, Leucine, and Methionine deficient Neurobasal media (Athenaes) supplemented with L-Arginine hydrochloride, L-Leucine, and L-Methionine were used during differentiation and neuron maintenance.

DIV2 motor neurons were lysed in volumes ranging from 600 to 1000 microliters of 8M Guanidinium Hydrocloride(GndHCl) and homogenized using a probe sonicator. Samples were centrifugated at a rcf of 14000g to remove solid debris, and protein concentration in the lysates was measured using a bicinchoninic acid protein assay kit (Micro BCA Protein Assay Kit, Thermo Scientific). Equal amounts (750 micrograms to 1 mg depending on the experiment) of protein labelled with D8 or 13C6 15N2 lysine (from wildtype and KO cells) were mixed, added phosphatase inhibitors (Sigma Phosphatase Inhibitor Cocktails 1 and 3) at 8-fold the recommended concentration, then treated with 8.8 mM DTT at 56°C for 15 minutes, followed by a 30-minute incubation at room temperature in the dark with 15 mM iodoacetamide. The samples were then diluted 8-fold with 100 mM ammonium bicarbonate to reduce GndHCl concentration to 1 M, and then added 2% (W/W) LysC (Wako). The samples were incubated 12 h at 37 °C. After that, another aliquot of LysC was added (2% W/W) and digested for additional 6 hours. After this, samples were acidified with formic acid to a final concentration of 5%. The digests were then desalted using a MAX-RP Sep Pak ® classic C18 cartridge (Waters) following the manufacturer’s protocol. Sep Pak eluates were dry evaporated in preparation for TiO2 based phophopeptide enrichment.

Phosphopeptide enrichment was performed in an AKTA Purifier (GE Healthcare, Piscataway, NJ) using 5 µm titanium dioxide (TiO2) beads (GL Sciences, Tokyo, Japan) in-house packed into a 2.0 mm x 2 cm analytical guard column (Upchurch Scientific, Oak Harbor, WA). Digests were resuspended in 240 µl buffer containing 35% MeCN, 200 mM NaCl, 0.4% TFA and loaded onto the TiO2 column at a flow rate of 2 ml/min. The column was then washed for 2 min with 35% acetonitrile (MeCN), 200 mM NaCl, 0.4% TFA to remove non phosphorylated peptides. Phosphopeptides were eluted from the column using 1 M Potassium Phosphate Monobasic (KH2PO4) at a flow rate of 0.5 ml/min for 30 min directly onto an on-line coupled C18 macrotrap peptide column (Michrom Bioresources, Auburn, CA). This column was washed with 5% MeCN, 0.1% TFA for 14 min and the adsorbed material was eluted in 400 µl of 50% MeCN, 0.1% TFA at a flow rate of 0.25 ml/min. The eluate was solvent evaporated and then resuspended in 240 µl 20 mM ammonium formiate pH 10.4 for fractionation of the peptide mixture by high pH RP chromatography. The phosphopeptides enriched sample was fractionated on an AKTA purifier system utilizing a Phenomenex Gemini 5u C18 110A 150 x 4.60 mm column, operating at a flow rate of 0.550 mL/min. Buffer A consisted of 20 mM ammonium formate (pH 10), and buffer B consisted of 20 mM ammonium formate in 90% acetonitrile (pH 10). Gradient details were as follows: 1 % to 9% B in 14 min, 9% B to 49% B in 4 min, 49% B to 70% B in 36 min, 70% B back down to 1% B in 3 min. 20 peptide-containing fractions were collected, evaporated and resuspended in 0.1% formic for mass spectrometry analysis.

Samples coming from RP fractionation were run onto a 2 mm, 75mm ID x 50 cm PepMap RSLC C18 EasySpray column (Thermo Scientific). 3-hour MeCN gradients (2–30% in 0.1% formic acid) were used to separate peptides, at a flow rate of 300 nl/min, for analysis in a Orbitrap Lumos Fusion (Thermo Scientific) in positive ion mode. MS spectra were acquired between 350 and 1250 m/z with a resolution of 500000. For each MS spectrum, multiply charged ions ions over the selected threshold (1E4) were selected for MSMS in cycles of 3 seconds with an isolation window of 1.6 m/z. Precursor ions were fragmented by HCD using a relative collision energy of 30%. MSMS spectra were acquired in the ion trap in centroid mode from m/z=110. A dynamic exclusion window was applied which prevented the same m/z (mass tolerance 60 ppm) from being selected for 30s after its acquisition.

### Peptide and protein identification and NeuCode quantitation

Peak lists were generated using PAVA in-house software^51^. All generated peak lists were searched against the human subset of the SwissProt database (SwissProt.2019.04.08), using Protein Prospector^52^ with the following parameters: Enzyme specificity was set as LysC, and up to 2 missed cleavages per peptide were allowed; Carbamidomethylation of cysteine residues was allowed as fixed modification; N-acetylation of the N-terminus of the protein, loss of protein N-terminal methionine, pyroglutamate formation from of peptide N-terminal glutamines, oxidation of methionine, 13C(6)15N(2) and 2H(8) labels in lysine, and phosphorylation on serine, threonine and tyrosine were allowed as variable modifications; Mass tolerance was 3 ppm in MS1 and 0.8 Da in MS2; The false positive rate was estimated by searching the data against a concatenated database which contains the original SwissProt database, as well as a version of each original entry where the sequence has been randomized; A 1% FDR was permitted at the protein and peptide level. For quantitation only unique peptides were considered; peptides common to several proteins were not used for quantitative analysis. Relative quantization of peptide abundances was performed by Protein Prospector, via integration of the MS1 ion intensities corresponding to the isotopic envelopes of K8 labeled species, in a ±30 second time window from MSMS identification.

### Data presentation and Statistical analysis

In most figures, data were represented as Mean ± standard deviation. In Sholl analysis figures of spot culture, data were represented as Mean ± standard error. Two-tail unpaired Student’s t-test or one way ANOVA were used using Prism (Graph Pad, La Jolla, CA, USA)

### Data Availability

The authors will make all data available to readers upon reasonable request.

---

**Figure S1.**
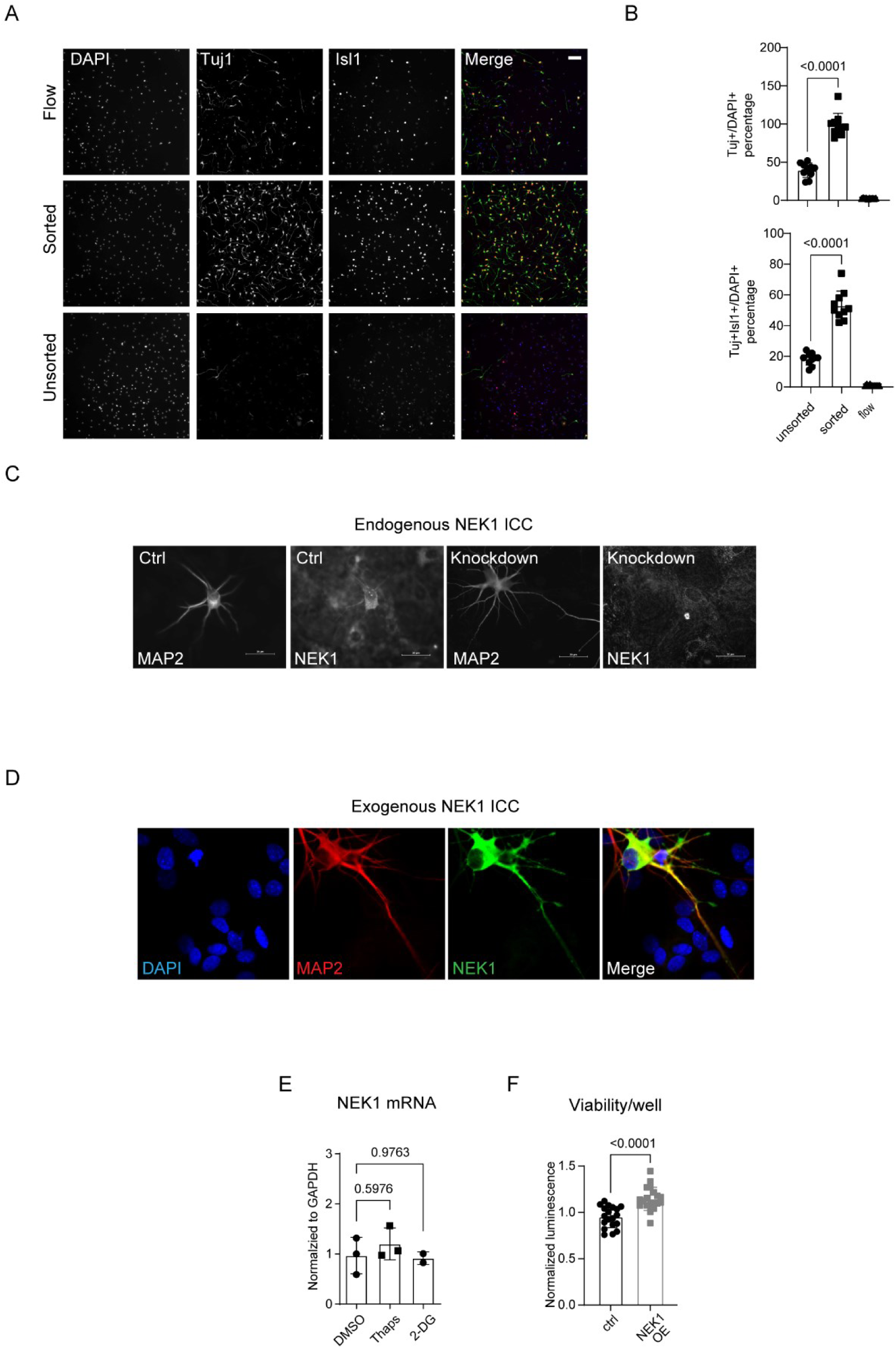
NEK1 expression in MN (related to figure 1) A. Representative images of differentiated cells before (unsorted) and after (flow and sorted) MACS purification. Cells are dissociated, purified, and seeded, then fixed one day later, stained for Tuj1 and Isl1, and counterstained for DAPI. Scale bar = 100 µm. B. Quantification of Tuj1^+^ neuron percentage and Isl1^+^ motoneuron percentage before and after MACS purification (n=10). C. Representative epifluorescence microscopy images of control and NEK1 knockdown motoneurons stained for MAP2 and NEK1. Scale bar = 20 µm. D. Representative confocal microscopy images of motoneurons transfected with human NEK1 cDNA and stained for MAP2 and NEK1. Scale bar = 20 µm. E. qRT-PCR analysis of NEK1 transcript expression in human motoneurons treated with thapsigargin (100 nM) and 2-deoxyglucose (2-DG) (1 mM) overnight (n= 2-3) F. Viability measurement of wildtype motoneuron overexpressing hsyn-NEK1 by CellTiterGlo assay (DIV=7) (n=19)

**Figure S2.**
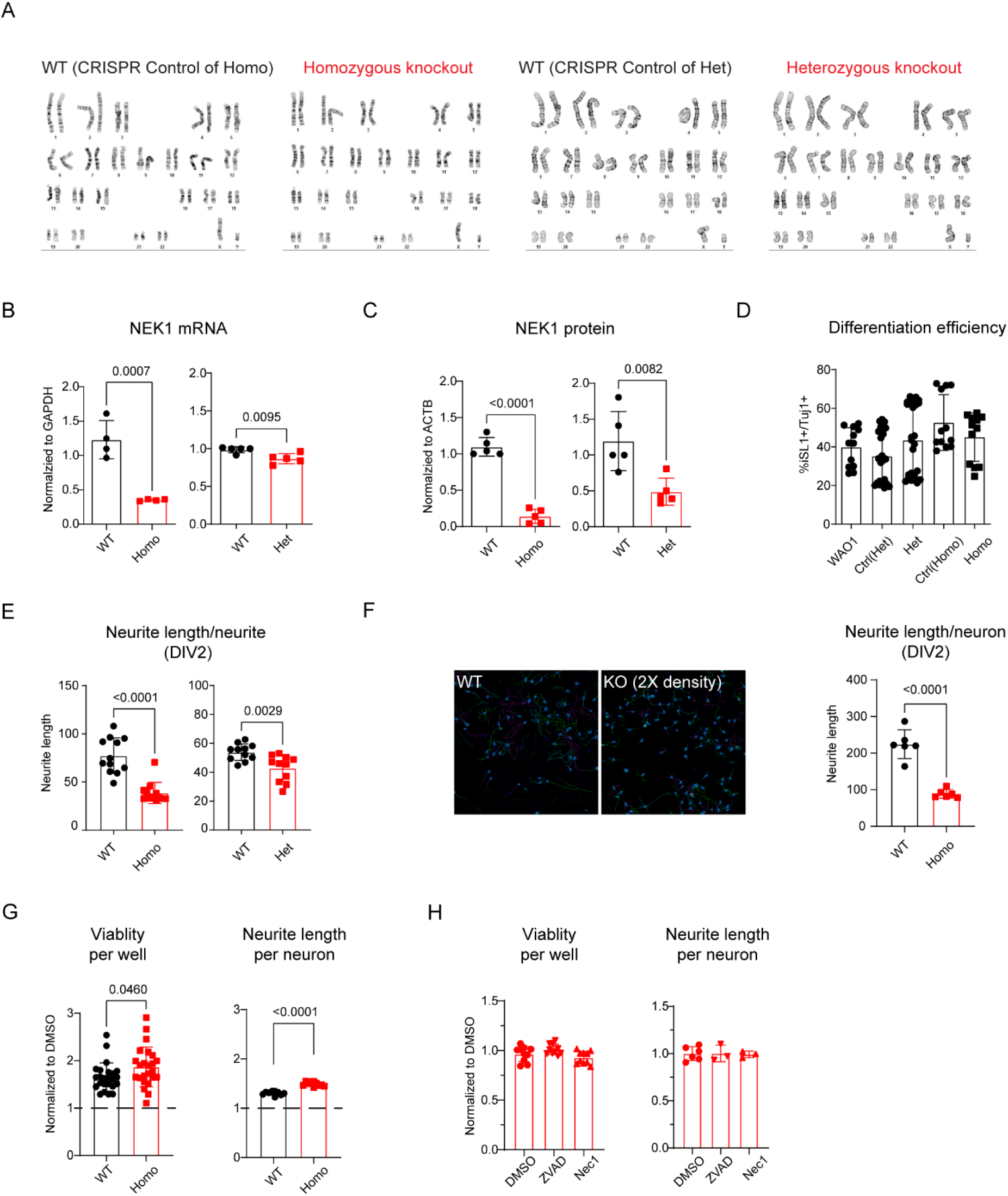
Loss of NEK1 leads to decreased neurite length (related to figure 2) A. Karyotype analysis of homozygous knockout (Homo), heterozygous knockout (Het) stem cells and their CRISPR unedited controls (WT) generated from the WAO1 ESC background to exclude unintended chromosome deletions caused by gene editing B. qRT-PCR analysis of NEK1 transcript in stem cells C. Western blot analysis of NEK1 protein in stem cells D. Quantification of Isl1^+^/Tuj1^+^ percentage of differentiated motor neurons (DIV=1) E. Quantification of neurite length per neurite (DIV=2) F. Representative images and quantification of wildtype and homozygous knockout motor neurons at DIV2. Homozygous knockouts were plated at twice the density of the wildtypes (n=6). G. Quantification of neurite length per neuron (n=20) and viability per well (n=10) after treating motor neurons with 20 µM blebbistatin for 48 hours (DIV2) H. Quantification of neurite length per neuron (n= 3-6) and viability per well (n=10) after treating *NEK1* knockout motor neurons with 10 µM Z-VAD or necrostatin-1 for 48 hours (DIV2)

**Figure S3.**
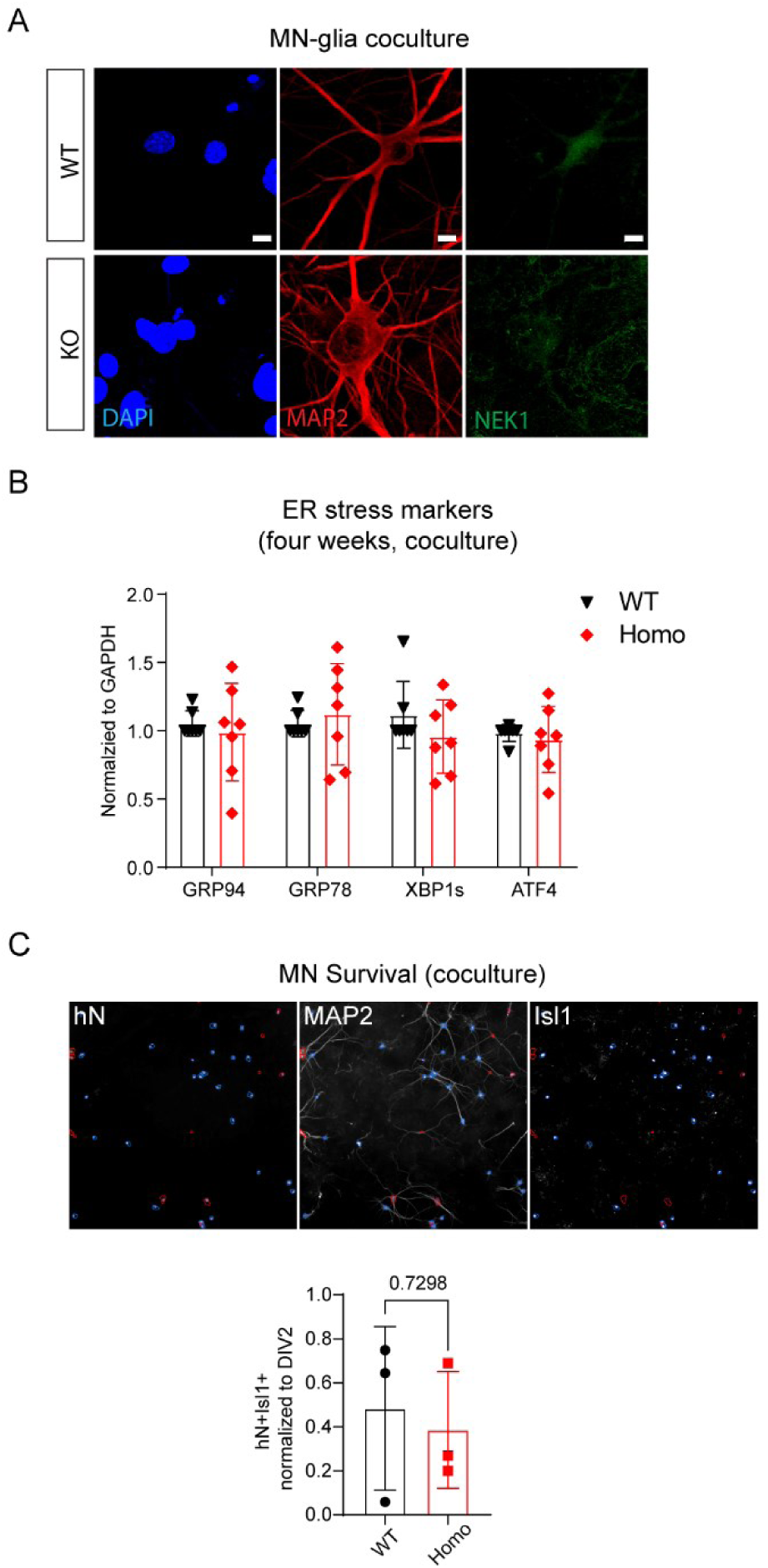
Glia is protective against NEK1 KO motor neuron death (related to Figure 3) A. Representative confocal images of four-week-old wildtype and homozygous knockout motor neurons cocultured with mouse glia, stained for MAP2 and NEK1, and counter stained with DAPI. Scale bar = 20 µm. B. qRT-PCR analysis of ER stress markers (human transcripts) from 4-week-old human motor neuron -mouse glia co-cultures (n=5-7) C. Representative epifluorescence images of 4-week-old motor neuron cocultures stained with human nucleus (hN), MAP2, and Isl1, with hN^+^Isl1^+^ cells selected in blue circles. The hN^+^Isl1^+^ neuron counts were normalized to the counts at DIV2 to evaluate cell death (n=3 different batches of coculture with >10 replica from each batch).

**Figure S4.**
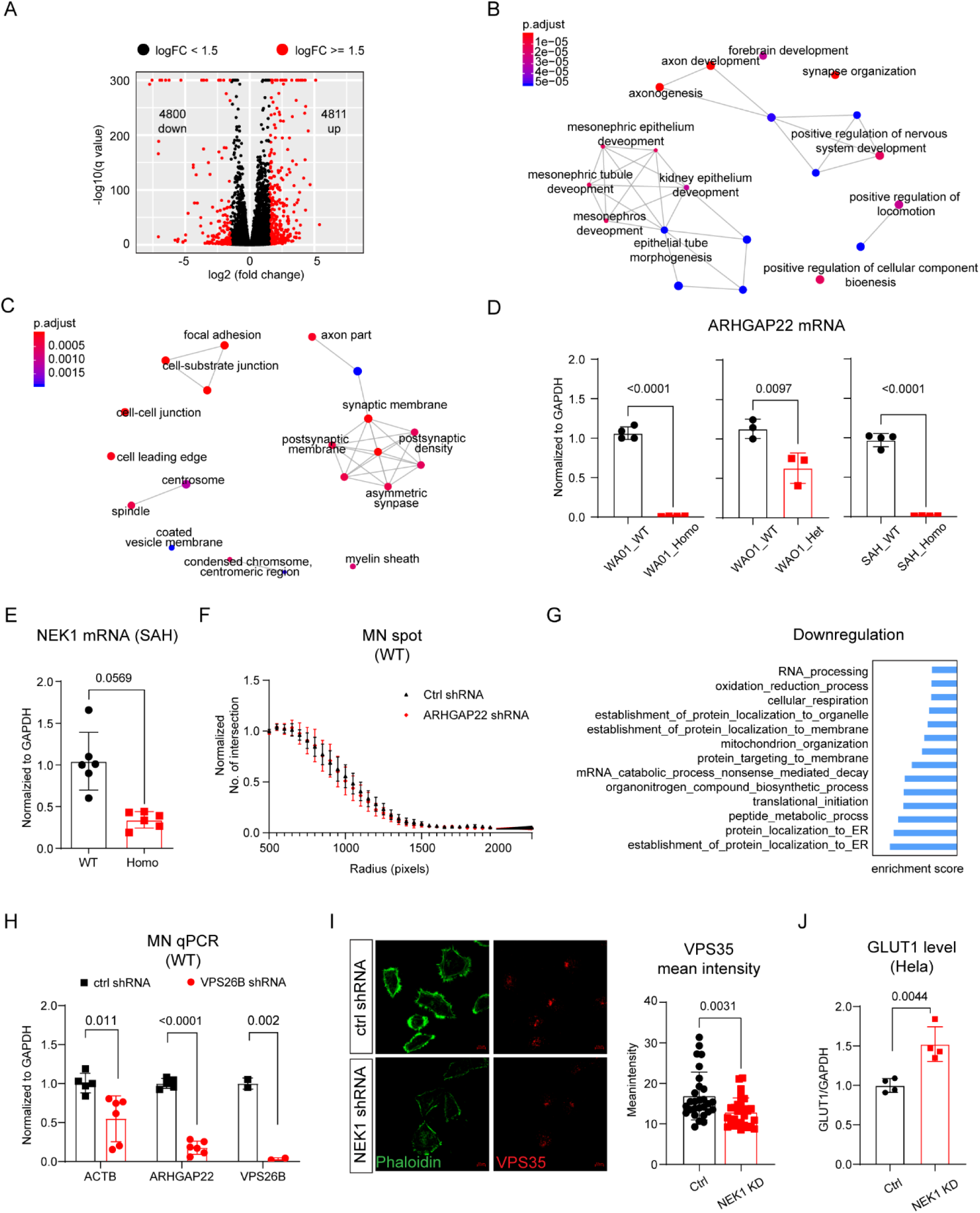
Loss of NEK1 dysregulates neurite outgrowth and protein transport related genes (related to Figure 4) A. Volcano plot showing transcripts with significant differential expression in DIV2 homozygous knockout MN (n=4) relative to wildtype controls (CRISPR unedited control n=4 and WAO1 control n=4) (FDR < 0.05) B. Network map showing the top 20 biological processes enriched in dysregulated transcripts (FDR < 0.05) in homozygous knockout MN (DIV2) C. Network map showing the top 20 cellular components enriched in dysregulated transcripts (FDR < 0.05) in homozygous knockout MN (DIV2) D. qRT-PCR analysis of ARHGAP22 in motor neurons (DIV2) E. qRT-PCR analysis of NEK1 transcript from wildtype and homozygous knockout stem cells generated in the SAH stem cell background F. Neurite outgrowth Sholl analysis of wildtype MN spot cultures treated with control or ARHGAP22 shRNA (n=7) G. Top down-regulated gene sets in heterozygous knockout MN by GSEA H. qRT-PCR analysis of ACTB, ARHGAP22, and VPS26B from wildtype motor neurons treated with control or VPS26B shRNA (n=5-6) for one week I. Representative confocal image of NEK1 knockdown Hela cells and the quantification. Cells are treated with control or NEK1 shRNA, stained for VPS35, counter stained with DAPI and Phaloidin-488. The mean VPS35 intensity is quantified. J. Quantification of GLUT1 expression in Hela cells measured by Western Blot. Hela cells are transfected with control or NEK1 shRNA construct, then the protein is collected four days after transfection.

**Figure S5.**
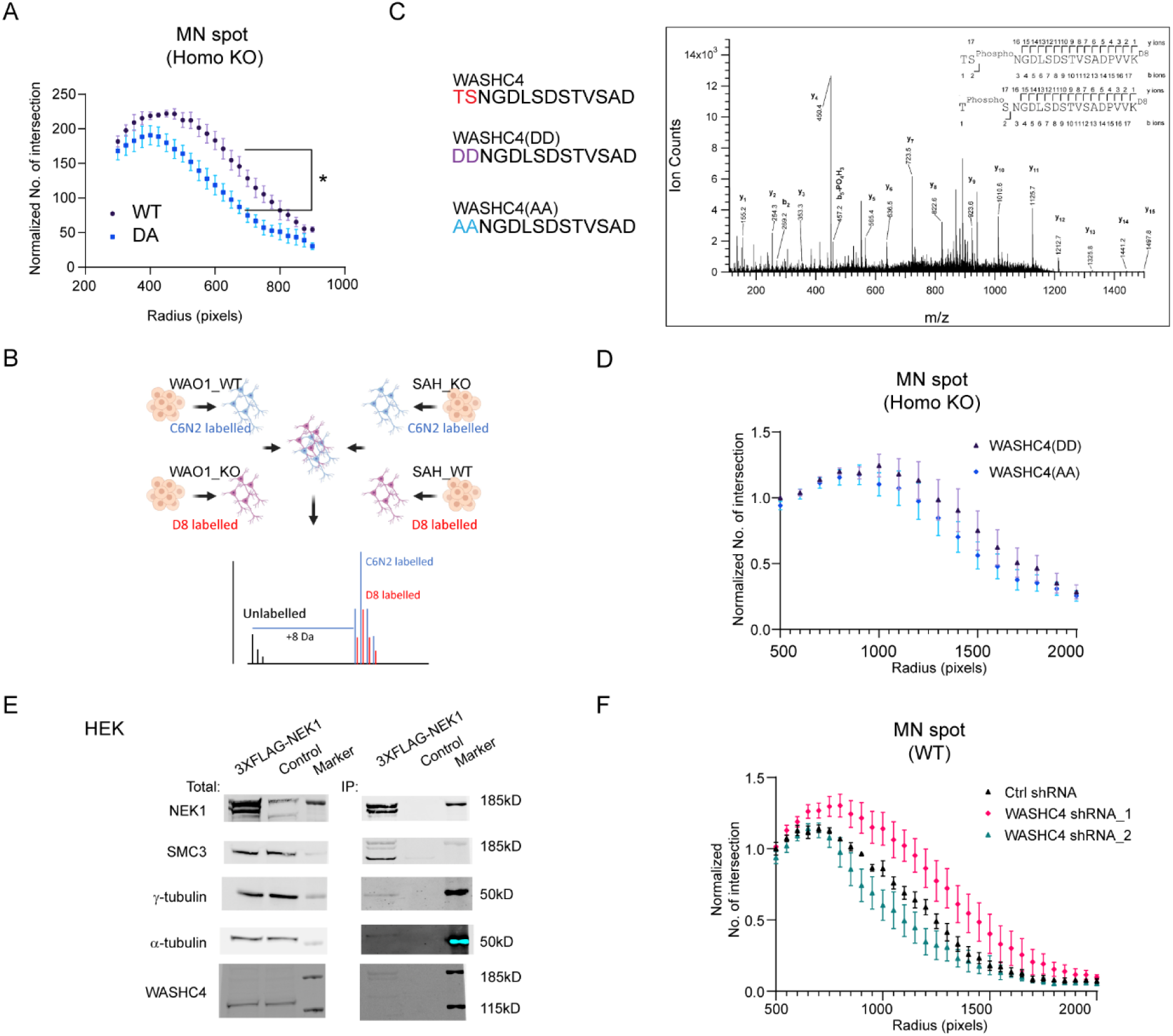
Loss of WASHC4 does not affect neurite length (related to Figure 5) A. Sholl analysis of neurite outgrowth of homozygous knockout MN spots overexpressing wildtype and D128A kinase dead mutant NEK1 driven by the human synapsin promoter (n=3 different batches of coculture with >10 replica from each batch) (DIV14) B. Illustration of NeuCode SILAC assay paradigm C. Left: WT and mutated sequences (phosphomimic S to D, and non phosphorylable S to A) around the T1156S1157 phosphorylation sites in SMC3. Right: MS spectra of the WASHC4 peptide. High-energy collision dissociation–tandem mass spectra obtained from precursor ion with mass 940.4432^+2^ found in LysC digests of D(8) (WT cells)/13C(6) 15N(2) (NEK1 KO cells)-Lysine labelled motoneurons, corresponding to the sequence T1156 to K1173 of human WASHC4 , phosphorylated at either T1156 or S1157 and lysine D(8) labelled. Masses of the N terminal sequence ion b_2_ show an increase of 80Da over the unmodified sequence, and no C terminal sequence ions (y) up to y_15_ are modified, limiting the location of the phosphate modified residue to the 2 first N terminal amino acids (T1156 or S1157). Observed sequence ions are labelled in the spectrum and marked in the sequence in the upper right corner. D. Neurite outgrowth Sholl analysis of homozygous knockout MN spots overexpressing phosphor-mimic WASHC3(DD) or phosphor-silent WASHC4(AA) mutants (n=9). E. Representative Western Blot images showing immunoprecipitation of SMC3 and tubulins with NEK1 in HEK cells overexpressing the 3XFLAG-NEK1 construct F. Neurite outgrowth Sholl analysis of wildtype MN spots treated with control or WASHC4 shRNA (n=4)

